# The Extracellular Loops of OmpA Control the Slow Rate of *In Vitro* Folding

**DOI:** 10.1101/2020.10.08.331546

**Authors:** Meghan W. Franklin, Jacqueline J. Stevens, Joanna Krise, Joanna S.G. Slusky

## Abstract

Outer membrane proteins are all beta barrels and these barrels have a variety of well-documented loop conformations. Here we test the effect of three different loop types on outer membrane protein A (OmpA) folding. We designed twelve 5-residue loops and experimentally tested the effect of replacing the long loops of outer membrane protein OmpA with the designed loops. Our studies succeeded in creating the smallest known outer membrane barrel. We find that significant changes in OmpA loops do not have a strong overall effect on OmpA folding. However, when decomposing folding into a fast rate and a slow rate we find that changes in loops strongly affect the slow rate of OmpA folding. Extracellular loop types with higher levels of hydrogen bonds had more instances of increasing the slow folding rate and extracellular loop types with low levels of hydrogen bonds had more instances of decreasing the slow folding rate. Having the slow rate affected by loop composition is consistent with the slow rate being associated with the insertion step of outer membrane protein folding.

## INTRODUCTION

Almost all proteins in the outer membranes of Gram negative bacteria have the same fold. They are β-barrels with each strand connected to the next strand via a loop-structure. The periplasmic loop-structures are known to be shorter and are traditionally referred to as ‘turns’. The extracellular loop-structures are longer and more varied and are referred to as ‘loops’. The barrel-shaped proteins of the outer membrane modulate the import, export, adhesion, and enzymatic function at the bacterial cell surface. Outer membrane proteins (OMPs) tend to fold rapidly into vesicles (Kleinschmidt, et al. 1999; Burgess, et al. 2008) but require the BAM complex to insert *in vivo* (Doerrler and Raetz 2005; Kim, et al. 2007).

The folding of outer membrane protein OmpA is particularly well studied. OmpA is an 8-strand, 4-loop barrel. Like other OMPs, OmpA can be folded in the presence of vesicles and its folding can be visualized and quantified with SDS PAGE. Because OmpA folding is so easily visualized many careful folding studies have been conducted to map out the OmpA folding pathway. These studies have illustrated the order of which parts of the protein contact the membrane (Kleinschmidt, et al. 1999; Kleinschmidt and Tamm 1999); the order of which parts of OmpA contact each other (Kleinschmidt, et al. 2011); and how folded different intermediates are (Danoff and Fleming 2017). Early work suggested that the loops contact the membrane first (Kleinschmidt, et al. 1999; Kleinschmidt and Tamm 1999) and that some hairpins were formed earlier than others (Kleinschmidt, et al. 2011).

Here we investigate the role of loops in OMP folding. Loops are the most difficult parts of proteins to accurately structurally predict or computationally design. Their greater range of φ and ψ angles and lower levels of homology among similar proteins make them the most difficult structure to predict (Gront, et al. 2012). Moreover, loops are often more dynamic than α-helices and β-sheets. However, the difficulty in designing loops or predicting their structure makes them no less important. Because loops are frequently observed on the surface of proteins, loops are often important in substrate recognition, receptor binding, and enzymatic activity in globular proteins (Rose, et al. 1985).

The loops of OMPs are often the functional part of the protein. Within OMPs the loops are known to control channel gating (Yildiz, et al. 2006; Zahn, et al. 2015), receptor binding (Fox, et al. 2014), and enzymatic activity (Kingma, et al. 2000; Vandeputte-Rutten, et al. 2001). Each OMP has half the number of loops as the number of β-strands in the barrel. For prototypical, single chain barrels, the number of β-strands per barrel vary from 8-26 (Franklin, Nepomnyachyi, et al. 2018). The most frequent number of amino acids per loop is five. These five-residue extracellular loops have five preferred geometries each with different position-specific amino acid preferences (Franklin and Slusky 2018). It is these five-residue loops which we design and assess for effects on OmpA folding.

Here we assessed the effects of drastically altering the loops of OmpA on protein folding. We use the previously described position-specific amino acid preferences of extracellular loop-types to design twelve loops and we test the effects of those loops on the rate of OmpA folding. Finally, we compare three methods of loop structure prediction for three canonical loop-types.

## RESULTS

The more frequently a given structure exists in nature the more designable it is believed to be. Our previous work determined that five-residue loops are the most common loop length and identified different types of five-residues loops (Franklin and Slusky 2018). For a five-residue loop the first and last positions are part of the previous and next β-strand, and the three middle residues constitute the connection between the strands as has been the convention since turns were first defined (Venkatachalam 1968). Because of the greater φψ variation in the central three amino acids, each loop was named based on the part of the Ramachandran map in which each of the three middle positions are localized as previously defined (North, et al. 2011) (Figure 1A), for example PLL has the three central residues in the proline, left, and left regions of the Ramachandran map. By definition, each loop type, such as PLD, AAL and PLL, has a different conformation with different φψ angles (Figure 1B)). Each loop also has particular amino acids associated with each position (Figure 1C-E) which were determined as previously described (Franklin and Slusky 2018). Notably, AAL frequently has intra-loop hydrogen bonding whereas PLL and PLD rarely have intra-loop hydrogen bonding (Franklin and Slusky 2018) (Figure 1F).

**Figure 1.**
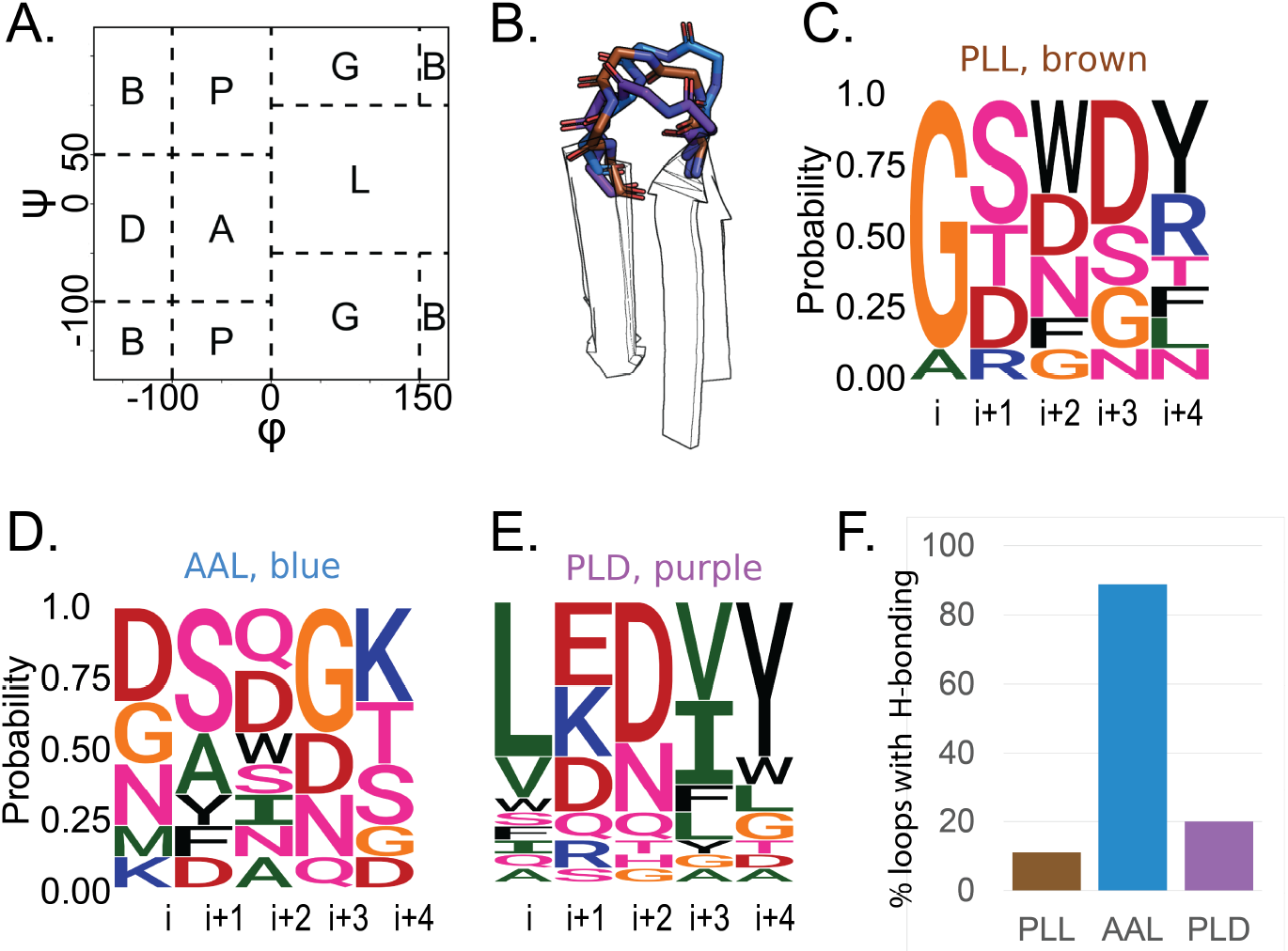
Loops with three letter nomenclature. **A)** The Ramachandran map divided up into five categories for describing backbone angles of loops. A is for alpha; B is for beta; G is for glycine; P is for proline; L is for loop. A five residue loop with backbone angles for the i+1 position in A, the i+2 position in A and the i+3 position in L would be called an AAL loop. **B)** Backbone trace of a representative loop for each of the three loop-types, shown in the assigned loop-type color (PLL, brown; AAL, blue; PLD, purple). **C-E)** Sequence logo plots showing the sequence diversity of each loop-type. The height of each letter represents the incidence of that amino acid at that position. **C)** PLL, **D)** AAL, **E)** PLD, **F)** Percent hydrogen bonding by loop type as previously determined. (Franklin and Slusky 2018)

### Designing Sequences for OmpA

In order to determine if shortening loops would affect OMP folding we redesigned each loop of OmpA with each of three loop types, PLD, AAL, and PLL. We chose to redesign the loops of OmpA because of how thoroughly OmpA folding has been characterized (Surrey and Jähnig 1992; Kleinschmidt, et al. 1999; Kleinschmidt and Tamm 1999, 2002; Kleinschmidt, et al. 2011; Danoff and Fleming 2017). The loops of OmpA are natively long, 17-21 residues in length for the NMR structure (Cierpicki, et al. 2006)(Figure 2A) and 10-20 residues in length for the crystal structure (Pautsch and Schulz 1998) (Figure 2B). Loop 1 and 4 are similar between the two structures and loops 2 and 3 are more different. Similar to most loops in our study of the loop types, the terminal positions of the long loops of OmpA fall into the β-region of the Ramachandran map because these residues are all strand by definition. These initial and terminal (i and I+4) φ and ψ angles are characteristic of two of the five types for five-residue loops. Loops 1, 3 and 4 have PLL-like terminal residues angles and loop 2 has AAL-like terminal residue angles (Figure 2C and 2D).

**Figure 2.**
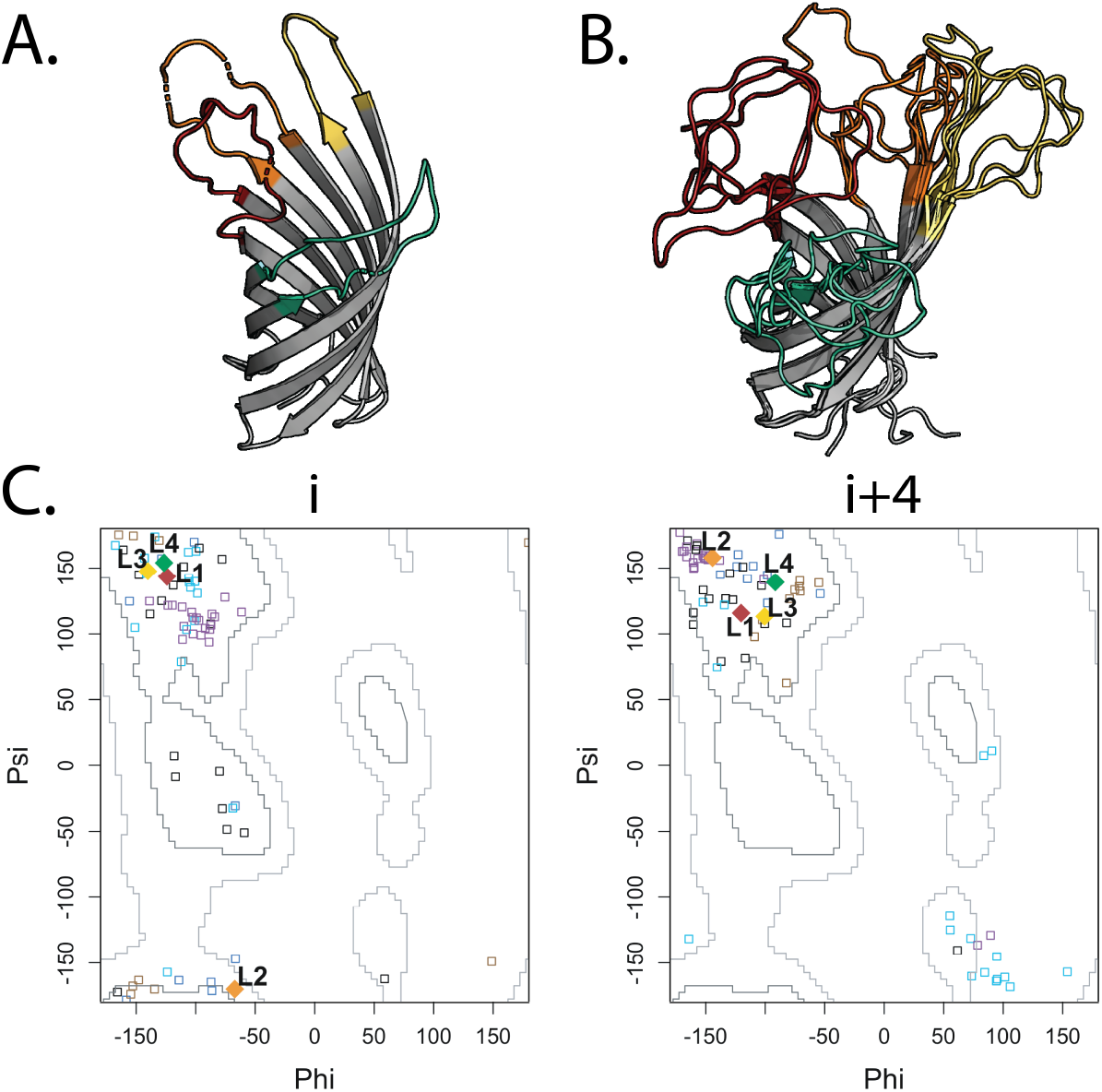
OmpA and its loop types. **A and B)** The four loops are colored red, orange, yellow and green from N to C-terminus. **A)** The loops of OmpA in crystal structure (PDB ID 1bxw) **B)** the first four states of the NMR structure (PDB ID: 2ge4). Some of the positions that are part of loops in the NMR structure are β-strands in the crystal structure.**C)** The φ and ψ angles of the i and i+4 positions of all the crystallized five-residue loops of OMBBs are shown in small empty squares, PLL in brown, PLD in purple, AAL in blue, and AAA in cyan. The native OmpA i and i+4 angles are shown in diamonds and colored red, orange, yellow, green for loops 1 to 4.

We built five-residue loops to substitute for the four naturally long loops of OmpA. Each loop position was substituted with 3 different sequences, one sequence with residues typical of an AAL loop, one sequence with residues typical of a PLL loop and one sequence with residues typical of a PLD loop. We used the position-specific amino acid preferences of each loop-type to select amino acids for the redesigned loops of OmpA (Figure 1 C-E). Twelve sequences in total were computationally selected from the three target loop-types to replace the four loops of OmpA (Table 1).

**Table 1.** Sequences designed for OmpA loops. Identity of the loops replaced and the 12 loop replacements.

| Loop ID | Residues | Native Sequence | PLL (Brown) | AAL (Blue) | PLD (Purple) |
| --- | --- | --- | --- | --- | --- |
| L1 | 38-56 | SQYHDTGFINNNGPTHENQ | GSFDT | DSNGK | WKNAA |
| L2 | 80-97 | YDFLGRMPYKGSVENGAYKAQ | GSFSR | KANDT | VDDVY |
| L3 | 125-140 | RADTKSNVYGKNHDTG | GRWDY | DSAGT | LKDIG |
| L4 | 164-183 | QFTNNIGDAHTIGTRPDNG | ATWSR | NSDGK | VSHGY |

### Folding Assay

DNA constructs of OmpA with designed loops were synthesized. Specifically, fifteen altered OmpA proteins were made, twelve were OmpA with one loop substituted and three were OmpA with all four loops substituted. The three constructs in which all four loops were substituted were substituted with loops of the same designed type (named by loop type, PLL-All Loops, PLD-All Loops, and AAL-All Loops). The fifteen proteins with designed loops and native OmpA were purified from inclusion bodies as described in the methods section. The folding of OMPs is assessable by using their property of heat modifiability. OMPs are so stably folded that a folded OMP shows a different migration on SDS-PAGE gel than when it is unfolded by boiling in SDS buffer (Rosenbusch 1974; Beher, et al. 1980).

Folding for these proteins can occur when they are presented with a synthetic membrane to fold into. To assess folding kinetics, each protein was rapidly diluted into folding buffer containing large unilamellar vesicles (LUVs). The protein LUV mix was incubated at 22°C while being shaken for up to six hours. Samples were taken at various time points, starting at sixty seconds. Folding was quenched with loading buffer. Samples were loaded and separated on a gradient SDS PAGE gel. Densitometry of the folded and unfolded bands were analyzed (Figure S1).

Percent folded protein was measured by dividing the density of the folded band by the sum of the folded and unfolded bands. Folding experiments were repeated in triplicate. For each experiment, the folding kinetics were best fit to a double exponential equation (supplemental equation 1). The folding of 14 of the 15 proteins fit to a two-state kinetic equation, which includes a value for the observed k_fast_ and k_slow_ (Figure 3). Only OmpA with all four loops substituted with AAL-type loops folded too quickly to be accurately measured using an SDS PAGE analysis and accurately fit to such a curve. The folding efficiency (fraction of protein folded at the final time point) for all except for PLL-All Loops are fairly similar to native OmpA with ~90% completion, however, PLL-All Loops only folded to ~61% (Figure 4A); similarly, most k_fast_ values were consistent with wild type OmpA ranging from 0.024-0.045 per second (Figure 4B). There was much greater variance the slow rate (Figure 4C), than the fast rate. The rate constant k_slow_ varied three orders of magnitude from ~1×10^−6^ to 4×10^−3^ per second. Wild type OmpA folding kinetics is consistent with previous measurements (Burgess, et al. 2008).

**Figure 3.**
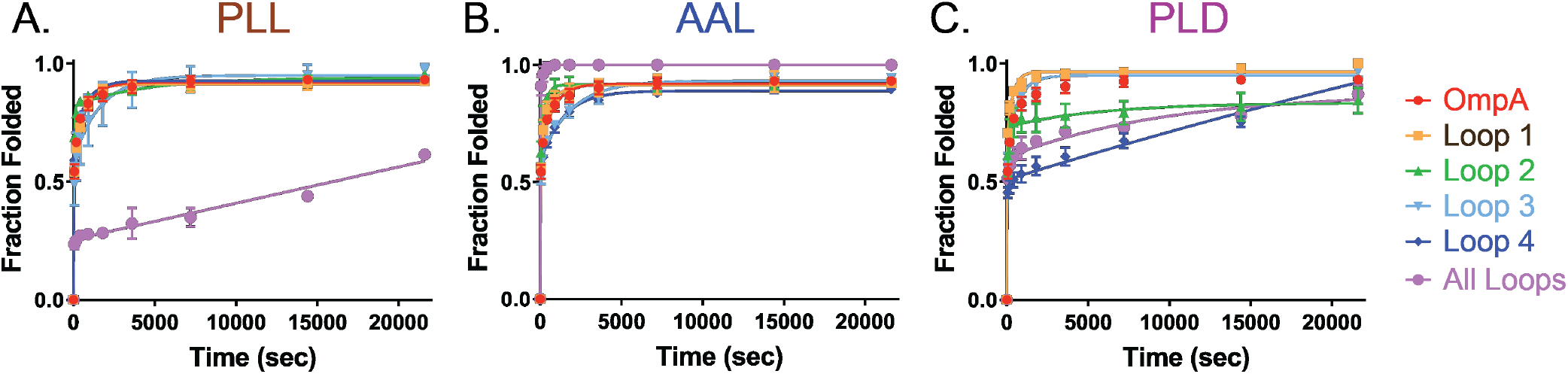
The fraction of OmpA folded over time at 22°C, determined by densitometry analysis of SDS-PAGE bands. Experiments were performed in triplicate with the average value shown and error bars corresponding to standard deviation; solid lines are the fitted exponential rate equation for each protein. Wild type OmpA is shown in red, mutation of the first OmpA loop to the designed sequence loop is shown in orange, mutation of second loop is shown in green, third loop is sky blue, fourth loop is royal blue, and all four loops mutated at once is shown in purple. OmpA loops have been substituted with designed **A)** PLL, **B)** AAL, and **C)** PLD loops.

**Figure 4.**
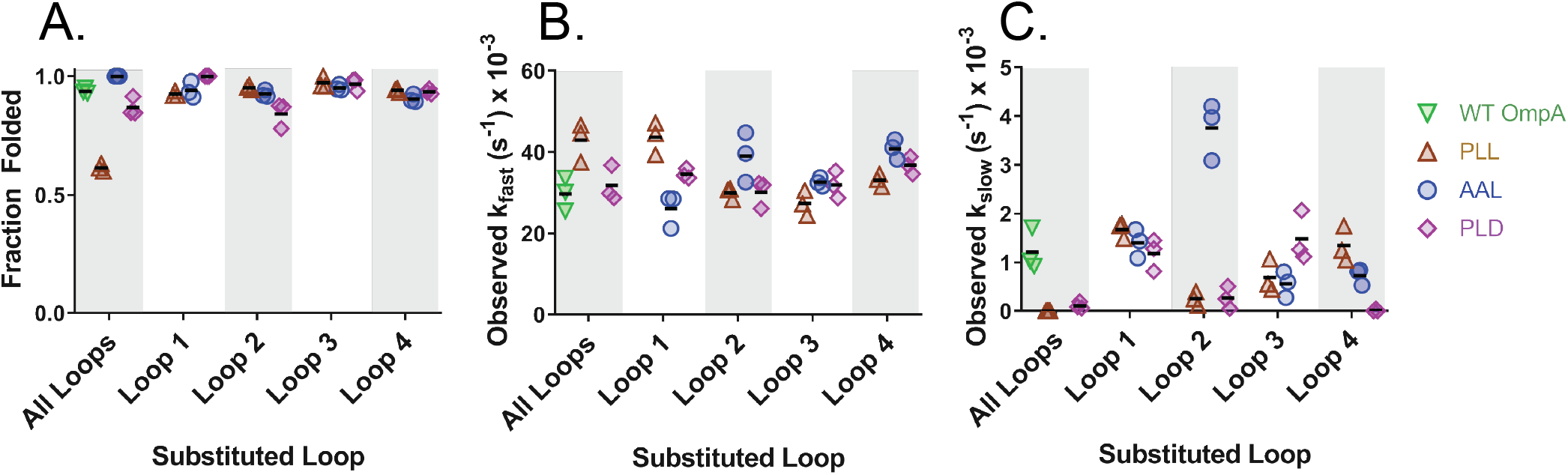
Analysis of kinetic data by substituted loop number. Each point represents the value obtained from each triplicate analysis, the average value is shown as a horizontal line. **A)** The folding efficiency of substituted loops at the end-point (6 hours). Wild type OmpA is shown in green, PLL loops are brown, PLD loops are purple, and AAL loops are blue. Observed **B)** k_fast_ and **C)** k_slow_ rate constants of OmpA with substituted loops are shown. Folding constants could not be shown for AAL-All Loops because it was too fast to accurately measure using this method. Values of the folding efficiency and rate constants in Table ST1.

Most loops tested were within a factor of two for both the k_fast_ and k_slow_. However, the distribution of the outliers was not uniform. The AAL-Loop 2 and AAL-All Loops both had a larger k_slow_, loop 2 by about three fold and all loops was too fast for measuring by this technique. In contrast, PLD-Loop 2 k_slow_ was five-fold lower, PLD-All Loops k_slow_ was ten-fold lower and PLD-Loop 4 k_slow_ loops was one hundred-fold lower than wild type OmpA. PLL- Loop 2 k_slow_ was five-fold lower and PLL-All Loops was fifty-fold lower than wild type OmpA.

### Structure prediction of outer membrane protein loops

Concurrently, we also assessed how accurately loop modeling programs could predict the structure of the three loop types we had experimentally tested. Though modern loop prediction software does not consider five-residue loops a challenge, we found consistent differences among the three loop prediction software packages that we used and we found that not only do AAL loops fold faster than the others, they are also easier to structurally predict than the other two loop types.

We predicted the loops of 72 native five-residue OMP loops that were previously typed as PLL, AAL, or PLD (Table ST2) (Franklin, Nepomnyachiy, et al. 2018). We predicted loop structure with PETALS (Wong, et al. 2017), DiSGro (Tang, et al. 2014) (Tang, et al. 2015), and the Rosetta KIC method (Mandell, et al. 2009). We rated performance by two metrics - backbone RMSD from native crystal structure (Figure 5 A-C, scatter plots) and similarity to native angles as measured by maintaining native loop type (recapture rates, Fig 5 A-C, bars). To illustrate the difference between measuring by RMSD and loop type angles, examples of models with low RMSD that did and did not recapture the native loop-types are shown in Fig 5D. We found significant differences among the three models in how well they modeled five-residue loops. For all three loop types, both of PETALS’ scoring functions – OSCAR-o and DBB - result in nearly identical performance as measured by RMSD (Fig 5, red and orange dots). Both functions also have 100% recapture of the AAL-type loops but result in different loop types other than native for the models that are of the PLL-type and PLD-type. The OSCAR-o function in PETALS recaptures the PLL loop-type better, while the DBB function recaptures the PLD type more effectively. mDiSGro has the next best performance based on RMSD (Fig 5, yellow dots). However, these models either recaptured the native loop-types or are not close to any loop-type (black bars). Finally, Rosetta KIC has the highest RMSD (Fig 5, green dots) and lowest recapture rates, often returning a model of other types instead of the native type.

**Figure 5.**
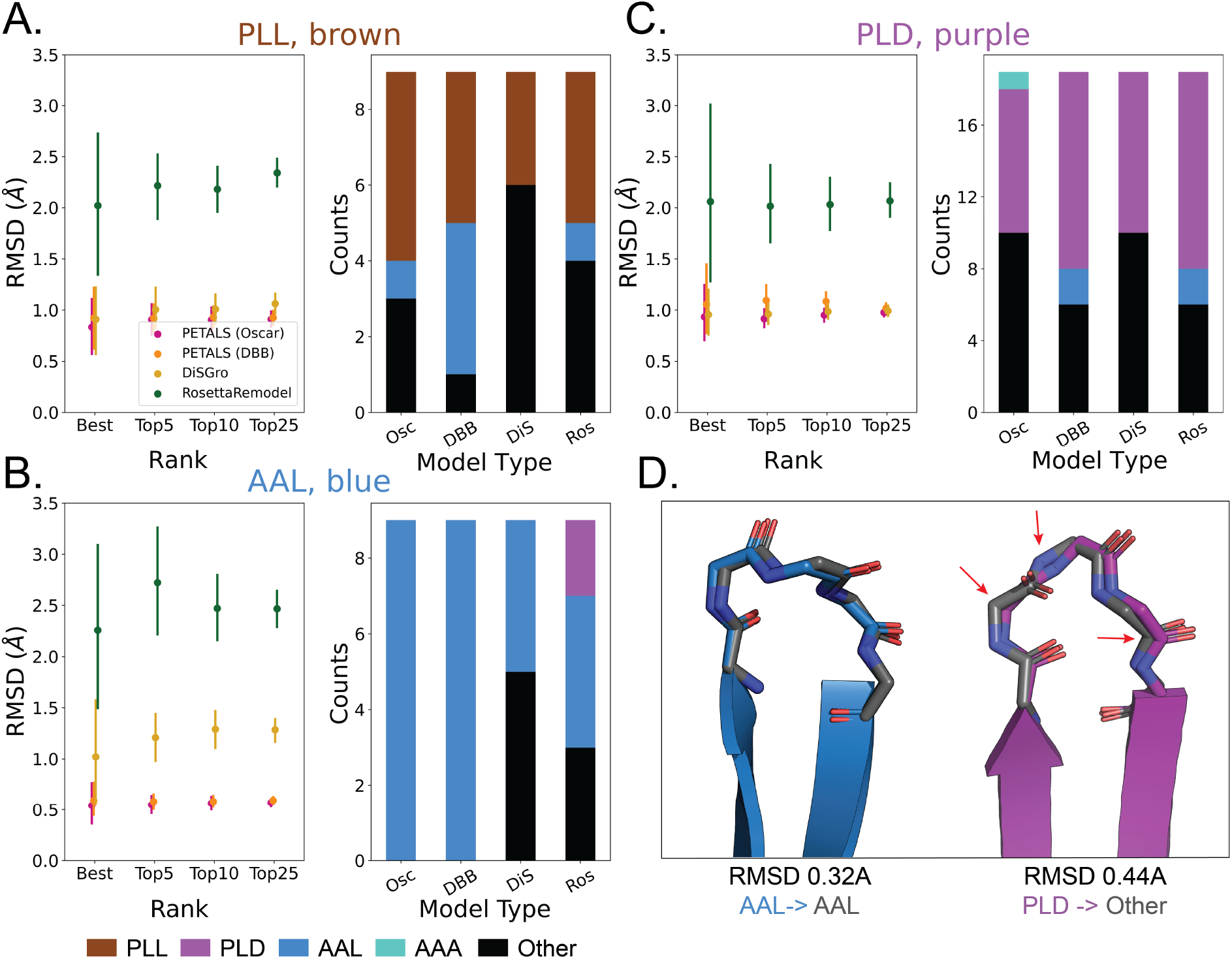
Testing loop modeling accuracy of three different modeling software. **A-C)** Average RMSD (left graphs) and loop-type recapture (right graphs) of 72 native five-residue OMP loops of three loop-types. The RMSD points represents the average backbone RMSD between the structure prediction and the native loop across the best designs, the top 5 designs, top 10 designs, or top 25 designs from left to right respectively. Lines represent 95% confidence interval. The bars represent whether the best model for each structure maintains all central angles (i+1 through i+3) within 30° of the loop-type cluster centroid; the expected color for all bars in a graph is the heading color. Black is outside every cluster; cyan is the AAA loop-type. The model types are PETALS (Oscar), PETALS (DBB), DiSGro, and Rosetta KIC from left to right. **A)** PLL, **B)** AAL, **C)** PLD. **D)** Lower RMSD models that do (left) or don’t (right) maintain loop type. The native loops are colored according to their native loop-type and the model is shown in grey. Both examples maintain a low RMSD, but the left example maintained the same loop-type assignment while the right example did not maintain the same loop-type assignment.

## DISCUSSION

Here we design 12 five-residue loops of three different types and test the effect of the 12 loops on folding. Overall there is not much change in the fast folding rates, but there is dramatic change in the slow folding rate. We find that the AAL loop-type, which is known to have more hydrogen bonding than other types is both easier to structurally predict and was, at least in some instances, able to achieve a much faster folding rate. Other loop types have slower slow folding rates.

### Loop folding

Though we made very large changes to the loops in OmpA we find that the changes in folding efficiency were minimal. Given the variety of loops it was surprising that almost all single loop mutants folded to completion similar to wild type. Although we found small differences, our overall finding was that folding efficiency remained markedly similar.

However, there was an overall difference in magnitude for how much the slow folding rate changed in comparison to how much the fast folding rate changed. While for every loop variant the fast folding rate was changed less than a factor of two, the slow folding rate was slowed by more than three orders of magnitude. We could also detect slow folding rate increases of at least four-fold (AAL- loop2). Moreover, one variant (AAL-All Loops) folded faster than could be accurately quantified using our SDS-PAGE methodology and would need other methods to get time points below 60 seconds to be accurately measured.

With respect to particular loop types, the AAL loops were both most predicted to correctly fold and they were the loops where a faster slow folding rate was seen. This is likely because the AAL loop-type is well positioned to have hydrogen bonding and is therefore a more energetically stable structure to make and to predict. AAL loops folded most consistently. PLD loops had the most folding variability. Finally, only the substitution of all four PLL loops substantially affected folding efficiency.

Although recent studies have shown that OmpA folding is most accurately modeled by using up to 11 states (Danoff and Fleming 2017), we have used the more usual single and double state kinetics for ease of comparison with previous studies from multiple labs over the past decades (Surrey and Jähnig 1995; Pocanschi, et al. 2006; Burgess, et al. 2008).

Many previous assessments and hypotheses around OMP folding into vesicles have started to converge to two stages of a folding pathway (Ricci and Silhavy 2019). The stages are roughly, pre-organization on the membrane and then insertion into the membrane. Because the loops are known to insert into the membrane first, we anticipate that the slow rate of folding which is most affected by changes in loops is the insertion into the membrane stage and that the fast rate of folding OMP folding is therefore most likely to be controlled by the formation of periplasmic turns and the resulting β-strand formation (Figure 5). This understanding is consistent with previous work that suggests it may be the turns and not the loops that facilitate the zipping up of OmpA strands (Danoff and Fleming 2017). However, we are agnostic about if the fast folding step is before (Figure 5A) or after (Figure 5B) the slow folding step as the hydrogen bonding of the β-strands may happen before or after insertion begins. Previous work shows β-strand formation late in the folding process indicating that hydrogen bonding of the β-strands is more likely to be part of the second step (Danoff and Fleming 2017). More work will need to be done to determine universality of this process for *in vitro* insertion of other outer membrane proteins.

**Figure 5.**
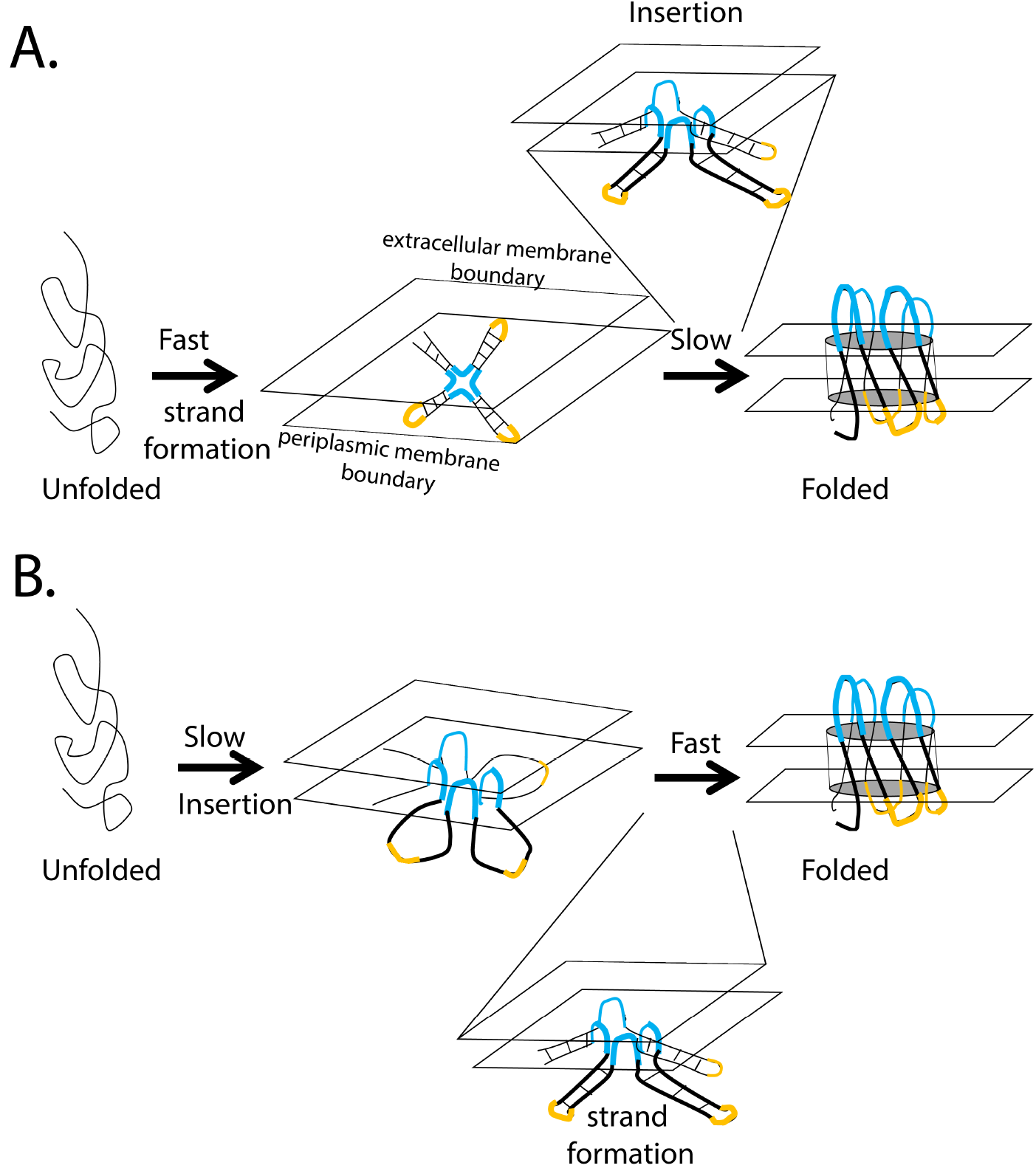
Two models of OmpA folding. Insertion is slow. Loops-region shown in blue, turns-region shown in yellow. **A)** Strand formation first and then insertion. **B)** Insertion first and then strand formation.

### Assessing loop prediction algorithms

The three methods used for loop structure prediction have some similarities and some differences. Both PETALS and DiSGro use energy functions to evaluate loops and both use the CSJD closure method (Coutsias, et al. 2003). PETALS grows loops from the N and C termini with the loop closing in the middle of the loop. DiSGro grows loops from the N-terminus and close. Rosetta’s KIC method also uses kinematics to close loops (from N to C) but in this method the loop closure is done many times with fewer energy filters and then the loops are optimized from the closed positions. mDiSGro and PETALS are advantageous for the speed at which they produce models. Rosetta KIC is ~1000 times slower but has the advantage of being integrated with the rest of Rosetta’s design algorithms.

Although 5-residue loops are generally not considered to be a challenge for loop prediction algorithms we found consistent differences among these algorithms. PETALS was the best on both metrics and to be especially good at recapturing AAL loops. PETALs may be better at OMP loop prediction because of its method of growing the loop from both the N and C terminus. AAL loops may be the easiest to recapture of the loop types used because it is the only one of the three loop types that has been found as a cluster within soluble protein loop types as well. Given that all of these structure prediction algorithms use knowledge-based potentials, all of the algorithms are likely to be biased towards building loops in globular proteins. This bias is likely due to the relative scarcity of membrane protein crystal structures and the documented differences in soluble loop angle preferences from those of OMPs (Franklin and Slusky 2018). Implementation of more membrane protein structures in the databases for loop construction algorithm would likely significantly improve their ability to effectively predict OMP loop conformations.

### Smallest barrel and utility for design

In a previous study (Koebnik 1999), Koebnik shortened what were at the time believed to be the loops of OmpA (Vogel and Jähnig 1986) to three-residue loops creating a 130-residue barrel. In that study, both the individual loops and the all loops substituted together were found to fold *in vivo*. Since then the NMR structure of OmpA (Arora, et al. 2001) was solved showing significantly longer loops and shorter strands than had been thought, suggesting that while Koebnik had meant to create three residue long loops he had inadvertently created loops between five and ten residues long. More recent evidence indicates that long loops such as these may be structured in the native outer membrane LPS though not in vesicles or micelles (Schubeis, et al. 2020).

The barrels of our three constructs with all four loops replaced are only 117 amino acids. Though these barrels were expressed along with the 154-residue C-terminal periplasmic domain of OmpA that folds independently (Ishida, et al. 2014), the barrels themselves are 13 residues smaller than the previously known smallest barrel. Our barrels are also smaller than the smallest known native barrel, PagP which has just 139-residue barrel. Though our barrels are small, the majority of them still have undiminished folding efficiency. Such small barrels may be useful as minimalist folding models of outer membrane proteins.

Finally, as previously described, the loops of outer membrane β-barrels are often responsible for the protein function. Knowing how robust OmpA folding is even with extreme loop manipulation may allow for planning redesigns of the loops with new functions, such as receptors or enzyme active sites, without concern for folding.

## CONCLUSION

Overall, we find that drastic changes in OmpA loops minimally effects OmpA folding. Through a careful analysis, we find small changes in the fast folding rate and more striking changes in the slow folding rate. Hydrogen bonding in loops may increase the slow folding rate and other changes decrease the slow folding rate. Overall these results are consistent with the slow folding rate being the rate of OmpA insertion into the membrane and not the β-strand formation.

## METHODS

### Loop Definitions

Our prior work described the amino acid preferences observed in the four-, five- and six-residue strand connectors of OMPs. The strand connectors in OMP crystal structures were clustered by the φ and ψ angles of the central residues to define the amino acid preferences of each position (Franklin and Slusky 2018). We use the same definitions outlined therein. Briefly, a loop is the extracellular strand connector between two adjacent strands, inclusive of the last residue of the first strand and the first residue of the second strand, while turns are periplasmic strand connectors. When describing the five-residue loops designed here they are numbered by convention, i to i+4; by definition, the i and i+4 residues are strand residues.

### Designed Loops

For each of the three loop-types that represented a spectrum of side-chain:backbone hydrogen bonding (Franklin and Slusky 2018) - PLL (brown), PLD (purple), and AAL (blue), sequences were generated from the amino acid preferences for each position. Specifically, for each position a script chose an amino acid based on the weighted probability of each amino acid identity for each position. There was no sequence selection beyond the random generation based on amino acid preference at each position. Only one sequence was generated for each loop-type and loop-position and that sequence was the one that was incorporated into OmpA.

### Cloning and Expression of OmpA Modified Loop proteins

A plasmid expressing mature OmpA without the 22 amino acid signal sequence in the pET303 vector was purchased from Genscript for cloning of the modified loop constructs. Primers were designed for the replacement of each of the 4 loops using Q5 polymerase (NEB) PCR reaction and a KLD enzyme mix (NEB). The resulting plasmids were transformed into chemically competent DH5α cells and sequences were confirmed by Sanger double stranded DNA sequencing. All confirmed plasmids were transformed into BL21(DE3) cells for expression. Transformed cells were grown in Terrific Broth (Millipore) with 0.04 % v/v glycerol at 37°C with shaking at 250 rpm to an optical density of 0.6 at 600 nm. Expression was induced by the addition of 1 mM Isopropyl β-D-1-thiogalactopyranoside (IPTG) and incubation at 25°C with shaking at 200 rpm for 18 hours. Cells were then harvested by centrifugation for 30 minutes at 4,700 g at 4°C. Pellets were re-suspended in 50 mM Tris pH 8.5, 0.1 mg/mL lysozyme and 1 μM phenylmethylsulfonyl fluoride (PMSF) and incubated for 30 minutes on ice before being sonicated for 5 minutes on 50% duty cycle (i.e. 250V) with 10 seconds off, 10 seconds on. Magnesium chloride (5 μM) and DNAse (5 μg/ml) were then added and incubation on ice continued another 30 minutes. Brij-L23 (Alfa Aesar) was added to a final concentration of 0.1 % v.v. Inclusion bodies were centrifuged at 4700 g for 30 minutes at 4°C then washed 2 times with 10 mM Tris, 2 mM EDTA, pH 8.2. The pellet was then re-suspended in wash buffer and divided into 2 tubes before a final centrifugation (4700 g, 30 min., 4°C). Supernatant was removed and the inclusion body pellets were stored at −20°C.

### Preparation of Large Unilamellar Vesicles (LUVs)

LUVs were prepared using the methods previously described (Burgess, et al. 2008). 1,2-didecanoyl-sn-glycero-3-phosphocholine (PC-di_C10_) (Avanti Polar Lipids) was dissolved in chloroform at 5.66 mg/mL then dried to a thin film in amber glass vials under a stream of nitrogen. The vials of lipid were then lyophilized overnight to remove residual solvent and then stored at −20°C until use. The lipid was then re-constituted in 2.66 mL of 20 mM borate buffer pH 10 to give a final concentration of 3.75 mM, then gently vortexed. LUVs were prepared by extruding the 3.75 mM lipids 15 times through a 0.1 μm filter using a mini-extruder (Avanti Polar Lipids) and stored at 4°C until use.

### Folding and SDS-PAGE

Inclusion body pellets were solubilized using 8 M urea, 10 mM borate pH 10, and 2 mM EDTA. After centrifugation at 18,000 g for 10 minutes the supernatant was filtered through a 0.45 μm syringe filter. Protein concentration was determined by measuring the absorbance at 280 nm and proteins were then diluted to 50 μM. Confirmation of the identity of the proteins was carried out using SDS-PAGE and MALDI-TOF mass spectrometry. Folding buffer was prepared to contain 3.2 mM LUVs, 1 M urea and 2 mM EDTA in borate buffer pH 10. Folding was initiated with a one in eleven dilution of 50 μM protein into folding buffer LUVs in an amber glass vial, for a final concentration of 4.5 μM protein, 1.6 M urea, 2.9 mM PC-diC_10_ LUVs, 9 mM borate pH 10, 1.8 mM EDTA. Folding samples were incubated at 22°C with shaking at 500 rpm in a thermal mixer block (Thermo Scientific) and time points were taken starting at 1 minute. At each time point, folding was quenched by taking 10 μL of sample and adding it to 10 μL of 2x Laemmli SDS gel-loading buffer without reducing agent (BioRad), vortexed and stored at 4°C until analysis. Unfolded protein in 8M urea, 10 mM borate pH 10 was prepared at 5 μM and loaded onto a 4-20% Mini-Protean TGX pre-cast gel (BioRad). 10 μL of each folding sample was subsequently loaded onto the same gel. After electrophoresis the gel was stained with Sypro Ruby overnight (chosen for its broad linear dynamic range). After de-staining in 10 % methanol, 7% acetic acid in water the gel was imaged using the BioRad Gel Doc.

### Data Analysis and Calculations

Using densitometry, the fraction folded was calculated by dividing the intensity of the folded band by the sum of the folded and unfolded bands. For each experiment, the folding kinetics were fit to both a single and double exponential equation in Graphpad Prism 7 and the best fit was chosen according to R-squared value (supplemental equation 1). k_fast_ and k_slow_ are the two rate constants expressed in inverse seconds. The reported values and error bars (standard deviation) represent the average of three independent experiments.

### Predicting loop structure

The center three residues of each native loop were modeled individually using the coordinates of i and i+4 positions as starting points in the context of the whole structure. The modeled structures were ranked according to the scores assigned by each program, and the top 25 models from each program were further analyzed. For each loop structure we independently generated an ensemble of 500 models using DiSGro as implemented in mDiSGro (Tang, et al. 2014; Tang, et al. 2015), PETALS (Wong, et al. 2017), and KIC in Rosetta using the remodel application (Mandell, et al. 2009). In the first round for each loop, knowledge of the angles of the i and i+4 residues were maintained while the coordinates of the central three residues (non-terminal residues) were predicted. For the second round of prediction and for the designed sequences, all loop coordinates were removed and the sequences were modeled from the loop-adjacent i-1 and i+5 residue angles known and the angles for i to i+4 unknown (Fig 2). Default parameters for each program were used to generate each ensemble, except as follows: in mDiSGro, the number of conformations was 10,000 and retained conformations was 1,000. In PETALS, the number of seeds was set to 5,000. Multiple scoring functions are implemented in PETALS and reported in the results; the default for loop building is DBB but each final model was rescored using several other scoring functions, so we also used the OSCAR-o scores for comparison. For KIC in Rosetta, the flag max_kic_build_attempts was set to 250; in the required blueprint file.

The modeled structures were ranked according to the scores assigned during loop building, and the top 25 models were further analyzed for RMSD and recapture rates to determine the best model. RMSD values were calculated for the backbone atoms of all amino acids in the loop structure. A loop model was determined to be “recaptured” if all the φ and ψ angles of the non-terminal amino acid positions were less than 30° from the native structure.

## ACKNOWLEDGMENTS

We gratefully acknowledge Jimmy Budiardjo, Rik Dhar, Ryan Feehan, Daniel Montezano, Paul Ikujuni, and Jaden Anderson for helpful conversations and NIH awards DP2GM128201, T32-GM008359, NSF MCB160205, The Gordon and Betty Moore Inventor Fellowship and KU-startup funds.

## Supplemental

**Figure S1.**
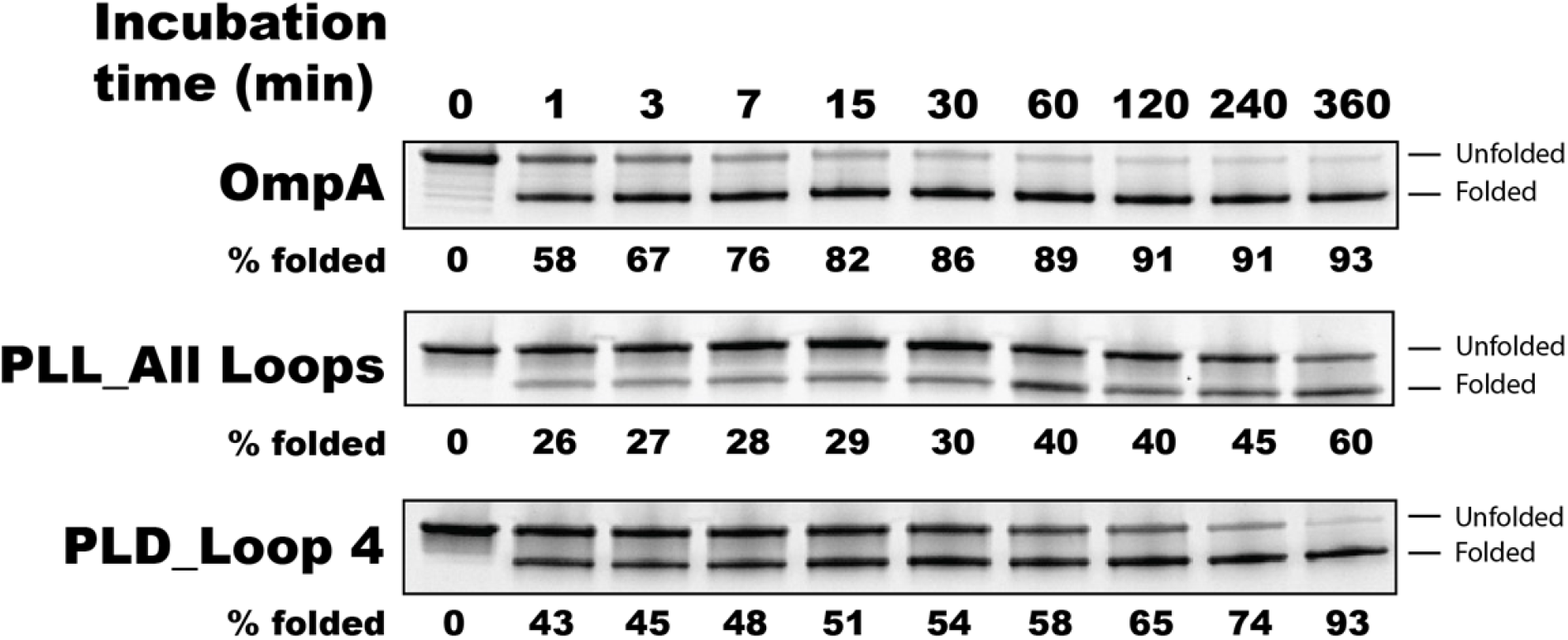
Representative de-stained SDS-Page gel of folding time course of substituted OmpA loops. Lane one is the unfolded protein in solubilization buffer (8 M urea) while lanes 2-10 are time points after mixing with LUVs in folding buffer. The folded protein runs faster than the unfolded protein. Data was collected in triplicate but only one gel of each is shown.

Densitometry analysis was performed and the amount of protein was calculated as Intensity_Folded_/(Intensity_Folded_ + Intensity_Unfolded_).

**ST1.** Tables of the values shown in figure 4. **A)** Fraction folded, **B)** k_fast_, **C)** k_slow_.

| A. | Fraction Folded |  |  |  |
| --- | --- | --- | --- | --- |
|  | OmpA WT | PLL | PLD | AAL |
| All Loops | 0.94 ± 0.01 | 0.62 ± 0.02 | 0.87 ± 0.04 | 1 ± 0 |
| Loop 1 |  | 0.93 ± 0.01 | 1 ± 0 | 0.94 ± 0.03 |
| Loop 2 |  | 0.95 ± 0.01 | 0.84 ± 0.05 | 0.93 ± 0.02 |
| Loop 3 |  | 0.97 ± 0.02 | 0.97 ± 0.03 | 0.95 ± 0.01 |
| Loop 4 |  | 0.94 ± 0.01 | 0.94 ± 0.01 | 0.91 ± 0.02 |
| B. | $k_{fast} \text{ sec}^{-1} \times 10^{-3}$ | | | |
|  | OmpA WT | PLL | PLD | AAL |
| All Loops | 29.75 ± 4.05 | 42.90 ± 4.78 | 31.80 ± 4.36 | N/A |
| Loop 1 |  | 43.66 ± 3.98 | 34.60 ± 1.23 | 26.1 ± 4.21 |
| Loop 2 |  | 30.00 ± 1.44 | 30.12 ± 3.48 | 39.03 ± 6.09 |
| Loop 3 |  | 27.38 ± 3.05 | 31.93 ± 3.33 | 32.60 ± 1.06 |
| Loop 4 |  | 33.13 ± 1.53 | 36.75 ± 2.11 | 40.82 ± 2.43 |
| C. | $k_{slow} \text{ sec}^{-1} \times 10^{-3}$ | | | |
|  | OmpA WT | PLL | PLD | AAL |
| All Loops | 1.22 ± 0.43 | 0.002 ± 0.003 | 0.11 ± 0.07 | N/A |
| Loop 1 |  | 1.68 ± 0.16 | 1.18 ± 0.33 | 1.4 ± 0.29 |
| Loop 2 |  | 0.25 ± 0.14 | 0.27 ± 0.23 | 3.76 ± 0.59 |
| Loop 3 |  | 0.69 ± 0.34 | 1.48 ± 0.51 | 0.56 ± 0.27 |
| Loop 4 |  | 1.35 ± 0.36 | 0.01 ± 0.02 | 0.73 ± 0.18 |

**Table ST2. List of all 5-residue loops.** (in excel file)

The folding kinetics were fit to a two-state association curve using a double exponential equation,

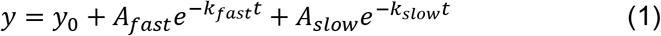

where *y* describes the fraction folded at time *t*. The fraction folded as time approaches infinity is described with *y_0_*, *k_fast_* and *k_slow_* are the rate constants and *A_fast_* and *A_slow_* are the negative amplitudes of each rate constant.

Other notes:

While our conditions (borate buffer instead of Tris, higher pH, lack of reducing agent, lower temperature) may not have been identical to other folding experiments for OmpA, our folding rate of k_fast_ = 29.7 × 10^−3^ and k_slow_ = 1.3 × 10^−3^ is similar to the published value of k_fast_ = 35 × 10^−3^ and k_slow_ = 3 × 10^−3^ (Burgess and Flemming, 2008).

## References

Arora A, Abildgaard F, Bushweller JH, Tamm LK. 2001. Structure of outer membrane protein A transmembrane domain by NMR spectroscopy. Nature Structural Biology 8:334–338.

Beher MG, Schnaitman CA, Pugsley AP. 1980. Major heat-modifiable outer membrane protein in gram-negative bacteria: comparison with the ompA protein of Escherichia coli. Journal of Bacteriology 143:906–913.

Burgess NK, Dao TP, Stanley AM, Fleming KG. 2008. β-Barrel Proteins That Reside in the Escherichia coli Outer Membrane in Vivo Demonstrate Varied Folding Behavior in Vitro. Journal of Biological Chemistry 283:26748–26758.

Cierpicki T, Liang B, Tamm LK, Bushweller JH. 2006. Increasing the Accuracy of Solution NMR Structures of Membrane Proteins by Application of Residual Dipolar Couplings. High-Resolution Structure of Outer Membrane Protein A. Journal of the American Chemical Society 128:6947–6951.

Coutsias EA, Seok C, Jacobson MP, Dill KA. 2003. A kinematic view of loop closure. Journal of Compuational Chemistry 25:510–528.

Danoff EJ, Fleming KG. 2017. Novel Kinetic Intermediates Populated along the Folding Pathway of the Transmembrane β-Barrel OmpA. Biochemistry 56:47–60.

Doerrler WT, Raetz CRH. 2005. Loss of Outer Membrane Proteins without Inhibition of Lipid Export in an Escherichia coli YaeT Mutant. Journal of Biological Chemistry 280:27679–27687.

Fox DA, Larsson P, Lo RH, Kroncke BM, Kasson PM, Columbus L. 2014. Structure of the Neisserial Outer Membrane Protein Opa60: Loop Flexibility Essential to Receptor Recognition and Bacterial Engulfment. Journal of the American Chemical Society 136:9938–9946.

Franklin MW, Nepomnyachiy S, Feehan R, Ben-Tal N, Kolodny R, Slusky JSG. 2018. Efflux Pumps Represent Possible Evolutionary Convergence onto the Beta Barrel Fold. Structure.

Franklin MW, Nepomnyachyi S, Feehan R, Ben-Tal N, Kolodny R, Slusky JSG. 2018. Evolutionary Pathways of Repeat Protein Topology in Bacterial Outer Membrane Proteins. eLife 7:e40308.

Franklin MW, Slusky JSG. 2018. Tight Turns of Outer Membrane Proteins: An Analysis of Sequence, Structure, and Hydrogen Bonding. J Mol Biol 430:3251–3265.

Gront D, Kmiecik S, Blaszczyk M, Ekonomiuk D, Koliński A. 2012. Optimization of protein models. Wiley Interdisciplinary Reviews: Computational Molecular Science 2:479–493.

Ishida H, Garcia-Herrero A, Vogel HJ. 2014. The periplasmic domain of Escherichia coli outer membrane protein A can undergo a localized temperature dependent structural transition. Biochimica et Biophysica Acta (BBA) - Biomembranes 1838:3014–3024.

Kim S, Malinverni JC, Sliz P, Silhavy TJ, Harrison SC, Kahne D. 2007. Structure and Function of an Essential Component of the Outer Membrane Protein Assembly Machine. Science 317:961–964.

Kingma RL, Fragiathaki M, Snijder HJ, Dijkstra BW, Verheij HM, Dekker N, Egmond MR. 2000. Unusual Catalytic Triad of Escherichia coli Outer Membrane Phospholipase A. Biochemistry 39:10017–10022.

Kleinschmidt JH, Bulieris PV, Qu J, Dogterom M, den Blaauwen T. 2011. Association of Neighboring β-Strands of Outer Membrane Protein A in Lipid Bilayers Revealed by Site-Directed Fluorescence Quenching. Journal of Molecular Biology 407:316–332.

Kleinschmidt JH, den Blaauwen T, Driessen AJM, Tamm LK. 1999. Outer Membrane Protein A of Escherichia coli Inserts and Folds into Lipid Bilayers by a Concerted Mechanism. Biochemistry 38:5006–5016.

Kleinschmidt JH, Tamm LK. 2002. Secondary and Tertiary Structure Formation of the β-Barrel Membrane Protein OmpA is Synchronized and Depends on Membrane Thickness. Journal of Molecular Biology 324:319–330.

Kleinschmidt JH, Tamm LK. 1999. Time-Resolved Distance Determination by Tryptophan Fluorescence Quenching: Probing Intermediates in Membrane Protein Folding†. Biochemistry 38:4996–5005.

Koebnik R. 1999. Structural and Functional Roles of the Surface-Exposed Loops of the beta - Barrel Membrane Protein OmpA from Escherichia coli. J. Bacteriol. 181:3688–3694.

Mandell DJ, Coutsias EA, Kortemme T. 2009. Sub-angstrom accuracy in protein loop reconstruction by robotics-inspired conformational sampling. Nature Methods 6:551–552.

North B, Lehmann A, Dunbrack RL. 2011. A New Clustering of Antibody CDR Loop Conformations. Journal of Molecular Biology 406:228–256.

Pautsch A, Schulz GE. 1998. Structure of the outer membrane protein A transmembrane domain. Nature Structural Biology 5:1013–1017.

Pocanschi CL, Patel GJ, Marsh D, Kleinschmidt JH. 2006. Curvature Elasticity and Refolding of OmpA in Large Unilamellar Vesicles. Biophysical Journal 91:L75–L77.

Ricci DP, Silhavy TJ. 2019. Outer Membrane Protein Insertion by the β-barrel Assembly Machine. EcoSal Plus 8:10.1128/ecosalplus.ESP-0035-2018.

Rose GD, Gierasch LM, Smith JA. 1985. Turns in peptides and proteins. Adv Protein Chem 37:1–109.

Rosenbusch JP. 1974. Characterization of the Major Envelope Protein from Escherichia coli: Regular Arrangement on the Peptidoglycan and Unusual Dodecyl Sulfate Binding. Journal of Biological Chemistry 249:8019–8029.

Schubeis T, Le Marchand T, Daday C, Kopec W, Tekwani Movellan K, Stanek J, Schwarzer TS, Castiglione K, de Groot BL, Pintacuda G, et al. 2020. A β-barrel for oil transport through lipid membranes: Dynamic NMR structures of AlkL. Proceedings of the National Academy of Sciences 117:21014–21021.

Surrey T, Jähnig F. 1995. Kinetics of Folding and Membrane Insertion of a β-Barrel Membrane Protein. Journal of Biological Chemistry 270:28199–28203.

Surrey T, Jähnig F. 1992. Refolding and oriented insertion of a membrane protein into a lipid bilayer. Proceedings of the National Academy of Sciences of the United States of America 89:7457–7461.

Tang K, Wong SWK, Liu JS, Zhang J, Liang J. 2015. Conformational sampling and structure prediction of multiple interacting loops in soluble and b-barrel membrane proteins using multi-loop distance-guided chain-growth Monte Carlo method. Bioinformatics 31:2646–2652.

Tang K, Zhang J, Liang J. 2014. Fast Protein Loop Sampling and Structure Prediction Using Distance-Guided Sequential Chain-Growth Monte Carlo Method. PLOS Computational Biology 10:e1003539.

Vandeputte-Rutten L, Kramer RA, Kroon J, Dekker N, Egmond MR, Gros P. 2001. Crystal structure of the outer membrane protease OmpT from Escherichia coli suggests a novel catalytic site. EMBO J 20:5033–5039.

Venkatachalam CM. 1968. Stereochemical Criteria for Polypeptides and Proteins. V. Conformation of a System of Three Linked Peptide Units. Biopolymers 6:1425–1436.

Vogel H, Jähnig F. 1986. Models for the structure of outer-membrane proteins of Escherichia coli derived from raman spectroscopy and prediction methods. Journal of Molecular Biology 190:191–199.

Wong SWK, Liu JS, Kou SC. 2017. Fast de novo discovery of low-energy protein loop conformations. Proteins: Structure, Function, and Bioinformatics 85:1402–1412.

Yildiz Ö, Vinothkumar KR, Goswami P, Kühlbrandt W. 2006. Structure of the monomeric outer‐membrane porin OmpG in the open and closed conformation. The EMBO Journal 25:3702–3713.

Zahn M, Bhamidimarri SP, Basle A, Winterhalter M, van den Berg B. 2015. Structural Insights into Outer Membrane Permeability of Acinetobacter baumannii. Structure 24:221–231.

